# Cholinergic Transmission in an Inducible Transgenic Mouse Model of Paroxysmal Dystonia

**DOI:** 10.1101/2024.03.15.585266

**Authors:** Mariangela Scarduzio, Karen L Eskow Jaunarajs, David G Standaert

**Affiliations:** Center for Neurodegeneration and Experimental Therapeutics; Department of Neurology, UAB, Birmingham, AL, USA

## Abstract

Altered interaction between striatonigral dopaminergic (DA) inputs and local acetylcholine (ACh) in striatum has long been hypothesized to play a central role in dystonia pathophysiology. Indeed, previous research across various genetic mouse models of human isolated dystonia has identified as a shared endophenotype with paradoxical excitation of striatal cholinergic interneurons (ChIs) activity in response to activation of dopamine D2 receptor (D2R). These mouse models lack a dystonic motor phenotype, which leaves a critical gap in comprehending the role of ACh transmission in the manifestation of dystonia. To tackle this question, we used a combination of *ex vivo* slice physiology and *in vivo* monitoring of striatal ACh dynamics in the inducible, phenotypically penetrant, transgenic mouse model of paroxysmal non-kinesigenic dyskinesia (PNKD). We found that, similarly to other genetic models, the PNKD mouse displays D2R-induced paradoxical excitation of ChI firing in *ex vivo* striatal brain slices. *In vivo*, caffeine triggers dystonic symptoms while reversing the D2R-mediated excitation of ChIs and desynchronizing the striatal cholinergic network. In WT littermate controls, caffeine stimulates spontaneous locomotion through a similar but reversed mechanism involving an excitatory switch of the D2R control of ChI activity, associated with enhanced cholinergic network synchronization. Together these observations suggest that D2Rs may play an important role in synchronizing the ChI network during heightened movement states. The “paradoxical excitation” described in dystonia models could represent a compensatory or protective mechanism that prevents manifestation of movement abnormalities and allows for phenotypic dystonia when lost.

## Introduction

Dystonia is a common movement disorder, affecting as many as 500,000 individuals in the United States alone [1]. Dystonia causes pain, disability, low quality of life, and sometimes death. Medication treatment strategies are of limited effectiveness and associated with adverse effects. Invasive methods, such as deep brain stimulation, are useful in some but not all cases. Though most commonly idiopathic, many genetic mutations have been shown to cause dystonia [2, 3]. Animal models that have mutations in several of these genes have been developed and offer new insights into its pathophysiology.

Dystonia is considered a network disorder, with the striatum playing a nodal role [4]. Typically, there is no evidence of neurodegeneration which points to the importance of events at the synaptic, neurochemical, and micro-structural level [5, 6]. Traditionally, proper control of movement has been attributed to antagonistic interactions between striatonigral dopaminergic (DA) inputs and local cholinergic (ACh) modulation in striatum [7]. Altered interaction between striatal ACh and DA has long been hypothesized to play a central role in dystonia pathophysiology [8, 9]. In support of this hypothesis, we have previously shown that three distinct mouse models expressing mutations in genes causing isolated dystonias (Dyt1ΔGAG/+, DYT1; Thap1C54Y/+, DYT6; Gnal+/-, DYT25) have enhanced activity (rather than the expected reduced activity) of cholinergic interneurons (ChIs) in response to dopamine D2 receptor (D2R) stimulation [10,11], a phenomenon termed “paradoxical excitation” [12]. The mechanisms underlying ChI paradoxical excitation in the genetic mouse models that we have studied to date involve an interaction between D2Rs and another G-coupled receptor (muscarinic receptors in Dyt1ΔGAG/+ and Thap1C54Y/+ and adenosine A2A receptor in Gnal+/-mice), which triggers a switch of D2Rs signaling from the canonical Gi/o to the non-canonical β-arrestin-mediated pathway [10, 11].

The central role of D2R modulation of striatal ChI activity in several aspects of striatal network functions, from synaptic plasticity formation to the control of motor- and reward-related behaviors has been extensively highlighted in recent work [13, 14, 15]. Precisely how the D2R dysfunctional control of striatal ChIs affects motor behavior in models of dystonia remains unexplored, largely because the most commonly studied models lack a recognizable dystonia phenotype. The exceptions are the mouse models for dopa-responsive dystonia (DRD) and Paroxysmal Non-Kinesigenic Dyskinesia (PNKD) [16, 17]. While the DRD model is difficult to utilize due to the developmental need for daily L-DOPA injections from the prenatal period, the PNKD model holds promise to dissect the striatal pathophysiology underlying dystonia, as indicated by recent studies utilizing this model [18].

PNKD is an autosomal dominant disorder caused by heterozygous mutations in the *PNKD* gene. Three isoforms of the PNKD protein are known; the longest one, PNKD-L, is expressed in neurons and found at presynaptic terminals where it regulates release of dopamine and other neurotransmitters [19]. Patients affected by PNKD develop attacks of dystonic posturing with some choreic and ballistic movements, often precipitated by coffee, alcohol, and stress [20]. The transgenic PNKD mouse model replicates many of these behavioral features, exhibiting a mix of inducible dystonia and dyskinesia upon exposure to stress or pharmacological treatments such as caffeine [16, 18]. The episodic and predictable nature of this response is particularly useful for understanding the physiological changes that occur during the onset of a dystonic attack. Indeed, recent work identified dysfunctions in striatal circuits of these mice associated with the expression of movement abnormalities [18]. However, the contribution of the cholinergic system has not yet been explored.

Recent work has shown that delta-rhythmic cellular dynamics in ChIs, which are reflected in spontaneous delta oscillations of ACh release [21], play an important role in regulating network rhythmicity and movement patterning [22]. These new observations were made possible by advances in techniques to track neurotransmitter release or neuronal membrane voltage *in vivo*. We have used this method of neurochemical reporter molecules and *in vivo* fiber photometry to investigate whether alterations in movement during inducible dystonic attacks in the PNKD mouse are associated with changes in striatal ACh rhythms *in vivo*. We also investigated the effects of the dystonia-inciting drug caffeine on ChI autonomous pace-making activity in *ex vivo* striatal slices. We found that in the PNKD mouse, caffeine induces involuntary movements with both dystonic and dyskinetic features in discrete episodes. The development of the motor phenotype is associated with a drastic reduction of the ACh delta rhythm. In contrast, WT mice show enhanced ACh delta rhythm in response to caffeine. These changes may arise from the interacting effects of caffeine and D2Rs in modulating ChI spontaneous firing.

## Material and Methods

### Mice

Male and female hemizygous mice (3-9 mos of age) of the PNKD-Tg strain developed by Dr. Louis Ptacek (B6.Cg-Tg(PNKD*A7V*A9V,-DsRed)704Ljp/J; JAX Stock #022146; Jackson Labs, ME, USA) were backcrossed with C57Bl/6J mice (Jackson Labs) for at least 5 generations, resulting in nearly congenic PNKD-Tg+/-(PNKD-Tg) and PNKD-Tg-/-(PNKD-WT) littermate controls. Traditional PCR genotyping was completed for each mouse used for breeding and experimentation with the primers as follows: Transgene detection, 1) 5’-ACATACATGCAGGAAAACATCC-3’, 2) 5’-AGTGCAGTCGTAAAAGTCAGAAC-3’; Internal Positive Control, 3) 5’-CTAGGCCACAGAATTGAAAGATCT-3’, 4) 5’-GTAGGTGGAAATTCTAGCATCATCC-3’. Data collection from different experimental groups was interleaved, mice were randomly assigned to groups, and experiments were blinded for all assays, including performing statistical analyses. All experimentation was approved by the University of Alabama Institutional Animal Care and Use Committee.

### Behavioral Assessment of PNKD Mice

#### Experimental Design

In a blinded, cross-over design, PNKD-Tg mice and their WT littermate controls (N=10-15/group), as well as a separate group of C57Bl/6J mice (N=4) never bred into the PNKD-Tg background, were randomly assigned to treatment groups. Mice were weighed and placed into a behavioral assessment cylinder (20×10 cm) placed on top of a white sheet of paper (to enhance visual contrast) for 1 h in order to habituate to the apparatus. Following habituation, mice were injected (10 mL/kg) with one of the following: Vehicle (0.9% saline), ±quinpirole HCl (1.0 mg/kg, ip), or caffeine (20 mg/kg, ip). All behavioral assessments were done live by a trained, blinded observer. Each animal was assessed at 10 min (or 20 min for microdialysis) intervals for 1 min each. Scores were recorded utilizing the Abnormal Involuntary Movement Scale (AIMs) for dyskinesia [23] and the Dystonia Disability Score (DSS) for dystonia [24]. Following assessment, mice were returned to the home cage. Different treatments were separated by a 3-7 d wash-out period.

#### Rating of Dyskinetic Movements

The AIMs is a well-established semi-quantitative scale useful for the assessment of the presence and severity of dyskinesia, as well as its anatomical manifestation [23]. In brief, each animal is assessed for 1 min for dyskinetic movements and postures in Axial (trunk, neck), Limb (hindlimb, forelimb), or Orolingual (face, mouth, tongue) planes. A score of 0-4 is given for each plane based on the following criteria: 0=no AIMs present for 1 min; 1=AIMs present for <30 s; 2=AIMs present for >30 s; 3=AIMs present for 1 min continuously, but interrupted by a tap on the behavioral apparatus; or 4=AIMs present for 1 min continuously and not interrupted by a tap on the behavioral apparatus.

#### Rating of Dystonic Movements

The Dystonia Disability Score (DSS) is an established semi-quantitative scale useful for the assessment of dystonia presence and severity, based on the inability of the animal to ambulate due to dystonic postures or movements [24, 25]. Each animal was assessed for 1 min every 10 min for dystonic disability and given a score of 0-5 based on the following criteria: 0=Normal behavior; 1=Inconsequential disability, slightly slowed or abnormal motor behavior; 2=Mild disability, limited ambulation unless disturbed, transient dystonic postures; 3=Moderate disability, limited ambulation even when disturbed, frequent dystonic postures; 4=Severe disability, almost no ambulation, sustained dystonic postures; 5=Maximal disability, prolonged immobility in dystonic postures. Importantly, a score of 1 or below indicates that dystonia is not present.

#### Assessment of specific abnormal movements

A behavioral inventory establishing the presence of specific abnormal movements was utilized to further define the nature and anatomical specificity of dystonic, dyskinetic, and otherwise abnormal movements. The inventory was similar to that used previously [26, 27]. Abnormal movements were assessed for 1 min every 10 min with their presence being denoted by a checkmark. Specific movements included tonic muscle activity (either flexion or extension), clonic muscle activity, twisting, and tremor, as well as overall changes in locomotor activity. Body area specificity was broken down into the following: face, neck, trunk, limbs, and tail. Behavioral scores were calculated by summing scores from each anatomical bin.

### *Ex vivo* Electrophysiological recordings of ChIs

Brain slices for electrophysiological recordings were obtained as previously described [10, 11]. Briefly, male and female PNKD mice (N=3/sex) and their control littermates (N=3/sex), 3–6 months of age were deeply anesthetized with isoflurane and perfused transcardially with ice-cold aCSF, bubbled with 95% O2-5% CO2, and containing the following (in mM): 2.5 KCl, 126 NaCl, 26 NaHCO3, 1.25 Na2HPO4, 2 CaCl2, 2 MgSO4 and 10 glucose. The brain was removed, blocked in the sagittal plane, and sectioned at a thickness of 280 µm in ice-cold aCSF. Slices were then submerged in room temperature aCSF bubbled with 95% O2–5% CO2 and stored at room temperature for at least 1 h before recording. The slices were transferred to a recording chamber mounted on an Olympus BX51WI upright, fixed-stage microscope and perfused (2 ml/ min) with oxygenated aCSF at 30°C. A 40X/0.9 numerical aperture water-immersion objective was used to examine the slice using standard infrared differential interference contrast video microscopy. Electrophysiological recordings were obtained with an Axopatch 200B amplifier (Molecular Devices), using borosilicate glass pipette pulled on a P1000 puller (Sutter Instruments). Patch pipette resistance was typically 3–4 MΩ when filled with aCSF. ChIs were identified by their distinct morphology (i.e., scarce, evenly distributed, very large neurons) and by the presence of spontaneous firing in the range of 0.5–6 Hz. ChI spontaneous activity was recorded in the loose, cell-attached configuration (seal resistance: 50 MΩ-200 MΩ), voltage-clamp mode, at a command potential at which the amplifier current is at 0 pA. Signals were digitized at 100 kHz and logged onto a personal computer with the Clampex 10.5 software (Molecular Devices). Spike frequency was averaged at 2 min intervals and normalized to baseline to yield the normalized event frequency, an indicator of how frequency changes over time and with drug treatments.

### *In vivo* Assessment of ACh Transients via Fiber Photometry

#### Experimental Design

PNKD (N=6) and WT (N=5) mice underwent a single stereotaxic surgery bilaterally in the dorsal striatum for viral injection and fiber optics implantation. Briefly, mice were anesthetized with 5% isoflurane and mounted in a stereotaxic frame. After exposing the skull, a craniotomy was performed at coordinates (from Bregma (mm) AP=+1; ML= +/-1.8) and AAV9-hSyn-GRABACh3.0 [28] (WZ Biosciences) was injected through a microinjector at DV= −3.2 mm from brain surface. Fiberoptic probes (200 µm in diameter, 0.37NA, 4mm long, Neurophotometrics, (NPM), CA) were implanted immediately after virus injection at the same injection location. Fibers were fixed on the skull using dental cement and two bone screws.Photometry experiments were carried out 25-50 days after the surgery. Mice were acclimated to the imaging arena for at least 30 min before simultaneously starting the acquisition of photometry data and video capture. PNKD and WT mice were imaged by placing them in an open field arena (acrylic 30×30 cm box) and allowing them to move freely while connected to a patch cord (200 µm in diameter, Doric). Photometry signals were acquired with the FP3002 system (NPM, CA) controlled via Bonsai software (Bonsai Foundation CIC), which also triggered the video camera. The green GRAB (GPCR-Activation-Based) ACh sensor was imaged with 470 nm LED and a 415 nm LED was used as interleaved isosbestic control to eliminate motion artifacts and photobleaching (i.e. ligand-independent signals). Both LEDs (light power adjusted to 50–100 µW) were alternatively turned on at 100 Hz. Data were collected for 15 min before and 45 min after treatment with vehicle (0.9% saline,ip), dystonia-inducing caffeine (20 mg/kg, ip) or quinpirole (1 mg/kg, ip) for a total of 60 min per session (**Figure 4A**). Each animal was recorded for only one session per treatment and had at least one rest day between sessions. The dose of caffeine was chosen based on preliminary experiments that allowed for maximum dystonia without seizure induction in >5% of animals. Daily for 7-10 days leading up to experiment day, mice received dummy injections while visiting the imaging arena. This habituation step drastically reduced stress-induced dystonic attacks in the PNKD mice.

#### Dystonia and Immobility Assessment

To assess dystonia during *in vivo* fiber photometry a time-based modified version of the DSS scale was used to take into account the duration of the disabilities. During each photometry session, a digital camera (J5create, JVCU100) was placed on top of the test cage and mice were video recorded for the duration of the experiment at a video frame rate of 30 frames/s. Videos were analyzed offline in 1 minute bins and severity scores were assigned by multiplying a 0-3 time-based grading scale, where 0=not present, 1=a given item is present for less than 30sec; 2=the item is present for more than 30 sec and 3=the item is present throughout the 1min monitoring period, by a 0-4 severity-based scale, where 0=no disability, 1=slow movement, 2=limited disability, 3=moderate disability and 4=severe disability, similarly to [29]. As quinpirole often inhibited voluntary movement in our preliminary experiments, we also assessed immobility during fiber photometry sessions. Immobility was defined as rigidity of the body and fixity of posture and was rated with a 0-3 time-based grading scale analogous to that used to rate dystonia.

#### Fiber Photometry Analysis

Video recordings of mice undergoing photometry were used to score immobility and dystonia as well as to video track the centroid of the mouse body to estimate the animal’s movement instantaneous speed using a custom-designed workflow in Bonsai software and post analysis with custom scripts in Matlab.

Photometry data were pre-processed using custom functions written in Matlab software (Matlab R2022a, MathWorks). In brief, mean fluorescence intensity values recorded from the 470 and 415 LEDs were deinterleaved. The isosbestic control data (415) was fit to the signal (least square linear regression) and the fitted control was used to calculate the change in fluorescence (ΔF/F) by subtracting and dividing the control data from the signal data. ΔF/F traces were transformed to the time-frequency domain computing Morlet-wavelets with the signal analysis toolbox for Matlab (*cwt* function, 18 voices per octave and frequency limits of 1-12 Hz). Power spectra averaged over genotypes are presented as colormaps. To quantify the change in power across different time windows within a single animal, the power spectrum was averaged over time in 10 min windows, then normalized to the total power as a sum across the frequency range 5.1–12 Hz, and further on represented as % of the total sum. To compare power only in the delta band between genotypes, the % of total sum power in each time window was averaged over 1.2 to 2.2 Hz frequency and then expressed as % of baseline change [30].

### *In vivo* Microdialysis of Striatal Extracellular ACh

#### Experimental Design

PNKD-Tg mice (N= 9) and their WT littermates (N= 7) were implanted with a microdialysis probe cannula (CMA7/2 mm, Harvard Apparatus) above the dorsal striatum, as previously [10, 11]. In vivo microdialysis for ACh was conducted 1-5 days after surgery, as previously described [10, 11]. Artificial cerebrospinal fluid (aCSF; 127.6mM NaCl, 4.02mM KCl, 750uM NaH2PO4, 2.1mM Na2HPO4, 2.00mM MgCl2, 1.71mM CaCl2; pH 7.4) was infused through a microdialysis probe (CMA7/2 mm, 7kDA-permissive membrane) at the constant flow rate of 2 uL/min while the animal was confined to an acrylic square chamber (30 cm W x 30 cm L x 50 cm H). The probe was implanted 2-4 h prior to sample collection. During habituation and continuing throughout the experiment, aCSF-containing Neo (100 nM to allow for reliable detection of the low levels of ACh) was perfused through a microdialysis probe (CMA7/2 mm, 7kDA-permissive membrane). Following habituation to the probe, baseline samples were collected at 20 min intervals for 1 h. After 1 h of baseline collection, mice were injected with Vehicle (0.9% saline, ip) and samples collected every 20 min for an additional 1 h, in order to correct for neurotransmitter response to the stress of injection. Mice were then injected with caffeine (20 mg/kg, ip) and samples collected for 2 h every 20 min. During baseline, vehicle-injection, and caffeine-injection sample collection, mice were assessed for dystonia, dyskinesia, and abnormal movement every 20 min. In a subset of mice, the D2R agonist, ±quinpirole (1.0 mg/kg, ip) was injected instead of caffeine with the same sampling parameters. At the end of sampling, mice were killed, and brains were removed and post-fixed with 4% paraformaldehyde. Fixed brains were then sectioned using a freezing microtome (Leica SM 200R, Buffalo Grove, IL) and cresyl violet staining was used to determine accurate probe placement. No animals were removed from the study due to improper probe placement.

#### HPLC-ED Analysis of Microdialysate

All microdialysis samples were analyzed for ACh using high-performance liquid chromatography with electrochemical detection (HPLC-ED; Eicom HTEC-500 with Eicom Autosampler INSIGHT, San Diego, CA) with an enzyme reactor (Eicom, AC-ENZYM II 1.0 x 4.0mm) and a platinum electrode (Eicom, WE-PT; +450 mV vs Ag/AgCl). The mobile phase consisted of 50 mM potassium bicarbonate, 134 µM ethylenediaminetetraacetic acid, and 1.22 mM decanesulfonate (pH 8.5) and was delivered at a rate of 150µL/min. Microdialysate sample chromatograms were analyzed based on established concentration curves for ACh (1-100 nM). The limit of detection was ∼10 fM.

### Statistical Analysis

Data are reported in text and figures as mean ± standard error of mean (SEM), with error bars in figures representing SEM (unless otherwise specified). Data were compared using GraphPad Prism 6 and 10 with the following statistical tests (as indicated in the figure captures): Student’s paired t-test for comparisons between paired data points, Student’s two-sample t-test or Mann-Whitney test for comparisons between un-paired data points, and 1-way or 2-way analysis of variance (ANOVA) followed by Sidak’s or Tukey’s Multiple Comparison Tests or otherwise specified for comparisons between multiple groups or different time points. For behavior, fiber photometry and microdialysis experiments N-values represent the number of mice. For electrophysiology recordings, n represents the number of ChIs and N the number of mice. Exact p values are provided in figure legends, and statistical significance in figures is presented as * p<0.05, ** p<0.01 and *** p<0.001. Though sex and age were included in the initial analysis as a factor, none of the parameters was altered by either variable and they were thus removed to simplify the data analysis.

## Results

### The Caffeine-Induced Phenotype of PNKD Mice Includes both Dystonia and Dyskinesia

Unstimulated, PNKD mice displayed few abnormal movements, typically the only notable feature was a wide-stance gait compared to their WT littermate counterparts. Minor abnormal postures and movements were induced by the stress of a vehicle injection in PNKD-Tg mice, reflected by increased scores compared to C57Bl/6J WT and PNKD-WT littermates on both the AIMs and DDS scales. These consisted of intermittent orolingual and limb dyskinesia, and splayed and/or hyperextended hindlimbs. This phenotype was drastically reduced by habituating the mice to the injection for at least 7 days before movement evaluation.

Caffeine administration induced hyperactivity and some stereotypy (paw shaking, repetitive cephalocaudal grooming, rearing) in PNKD-WTs. In PNKD-Tg mice, caffeine strongly induced bilateral dyskinesia and dystonia beginning approximately 10-20 min following systemic injection (**Figure 1A-B**). Dyskinesia was mostly manifested as limb and orolingual dyskinesia, particularly repetitive cephalocaudal grooming that was limited to the first and second steps (paw licking, followed by head grooming with the paws) (**Video 1**). Dystonic postures and movements connected to the limbs were prevalent, particularly sustained extension of the hindlimbs and hyperextended digits (**Figure 1C**, **Video 1**). A subset of mice also exhibited truncal hyperflexion, characterized by rearing up on the side of the cylinder with splayed hindlimbs and a concave back, tail hyperextension and brief head-shoveling movements (**Figure 1C**, **Video 1).** Despite enhanced abnormal movement, overall locomotion was depressed in PNKD-Tg mice treated with caffeine, compared to vehicle. A percentage of PNKD-Tg mice (<10%) did not display the dystonic/dyskinetic phenotype upon exposure to caffeine but, similarly to WT mice, they showed enhanced locomotor activity. In contrast, the effects the dopamine D2R agonists, quinpirole, on dystonic/dyskinetic behaviors were less consistent and a more prominent and generalized inhibition of activity was detected (**Suppl. Figure 2**).

**Figure 1.**
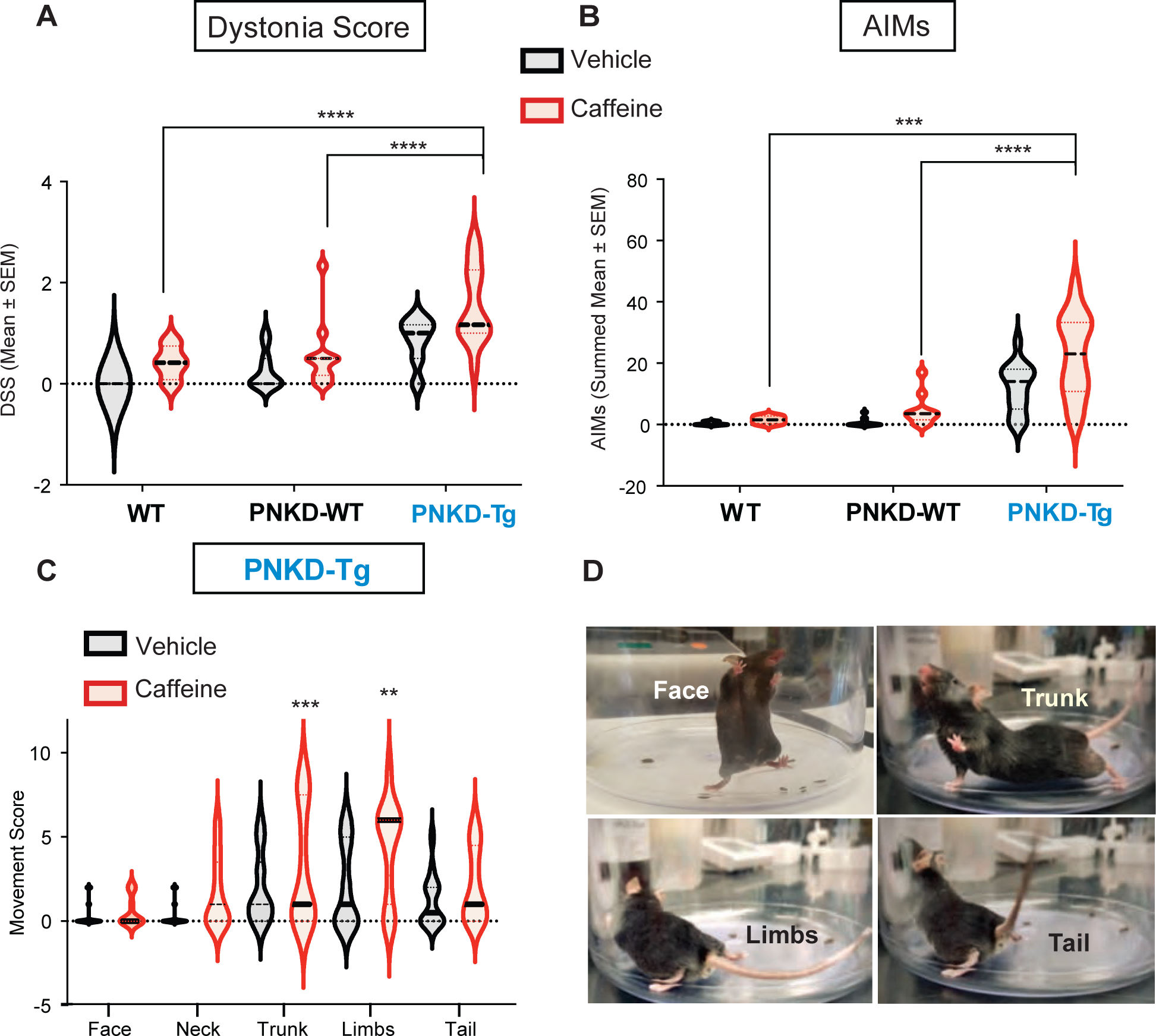
Dystonia and Dyskinesia are Inducible Components of PNKD Motor Dysfunction. Caffeine (20 mg/kg, ip) induced increases in both **A)** dystonia severity score (DSS, RM 2-way ANOVA, time: F_(2,74)_= 14.5, p<0.0001; genotype: F_(2,74)_= 34.4, p<0.0001; interaction: F_(4,74)_ =2, p=0.1; ** Tukey’s multiple comparisons test) and **B)** abnormal involuntary movements (AIMs, RM 2-way ANOVA, time: F_(2,74)_= 2.16, p=0.12; genotype: F_(2,74)_= 42.03, p<0.0001; interaction: F_(4,74)_ =1.02, p=0.4; ** Tukey’s multiple comparisons test) in PNKD-Tg vs WT and PNKD-WT mice over a period of 60 min with scores rated every 10 min for 1 min. **C)** Caffeine-induced dystonic movements in the PNKD mice interested especially the hind limbs, trunk and tail (2-way ANOVA, interaction: F_(4,130)_ =2.29, p=0.06, treatment: F_(1,130)_= 28.18, p<0.0001; body part: F_(4,130)_= 8.49, p<0.0001; ** Sidak’s multiple comparisons test) (**D)** Representative photos of the body parts interested.

### Caffeine and Quinpirole Have Opposing and Interacting Effects in Modulating Spontaneous Firing of Striatal ChI

To investigate whether behavior-inducing agents alter ChI activity at the cellular level, we recorded the spontaneous firing of dorsolateral ChIs in *ex vivo* striatal brain slices from PNKD mice and their WT littermates before and after acute application of caffeine or quinpirole **(Figure 2A)**. We found that basal firing rates of ChIs were no different based on genotype (**Figure 2B**). Bath application of quinpirole (10µM) produced a significant increase of ChI discharge in PNKD mice (**Figure 2C**), whereas little change in firing rate was seen in littermate controls (**Figure 2C**). Indeed, upon quinpirole treatment, PNKD mice showed significantly elevated ChI firing rates compared to littermate controls at the final time points, resembling the paradoxical excitation of ChI spontaneous firing described previously in non-phenotypic genetic models of isolated dystonia [10, 11].

**Figure 2.**
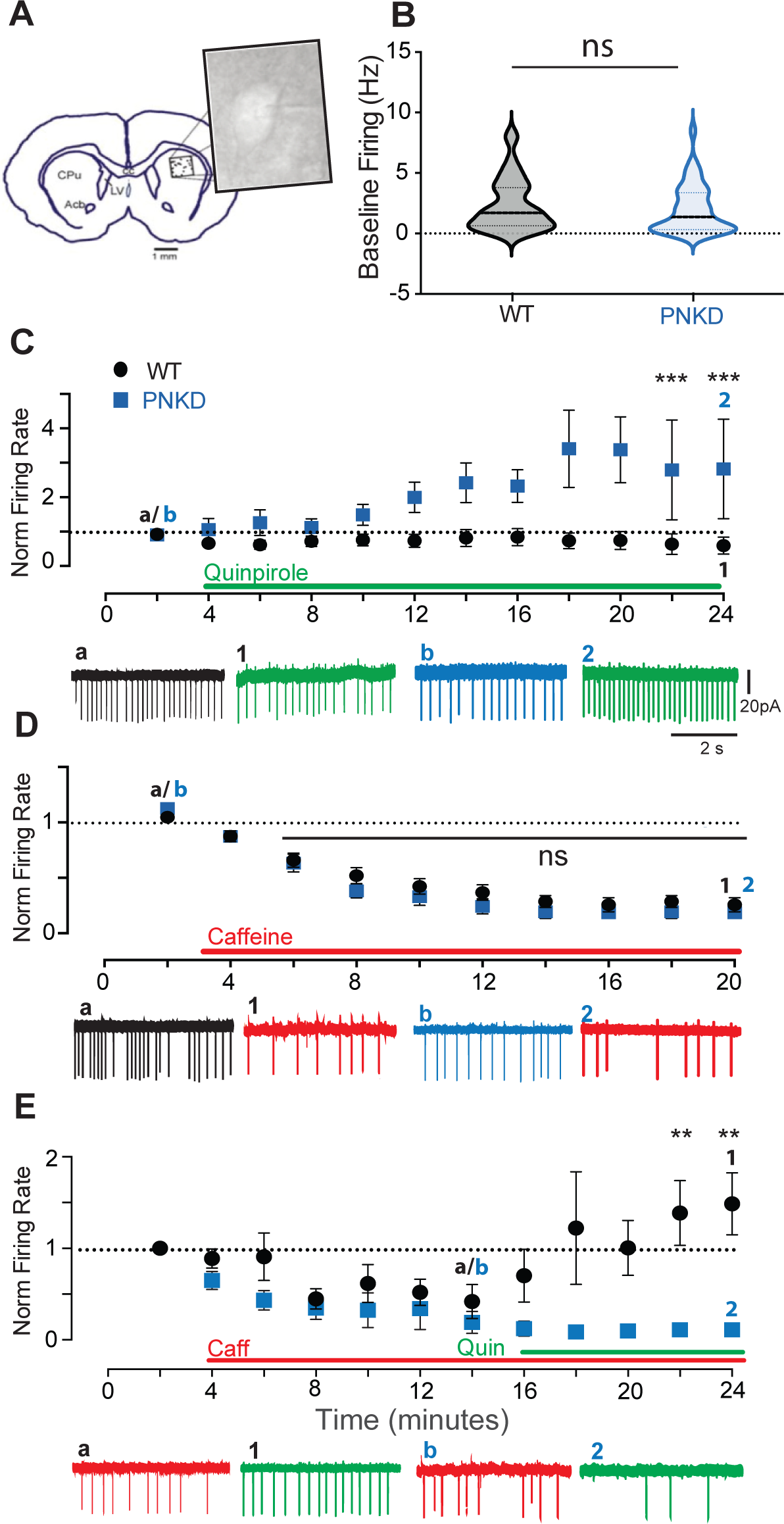
Caffeine and Quinpirole have opposing and interacting effects on ChI spontaneous firing. **A)** Experimental setting showing patch-clamp cell-attached recordings of striatal ChI in *ex vivo* striatal slices. **B)** Violin plot shows the summary statistics (line at mean +SEM) calculated from the mean ChI firing rate across all the neurons recorded at baseline from each animal: WT N=13, n=36; PNKD N=16, n=42, Mann-Whitney test, p= 0.47. Time courses and representative traces of the firing rate of ChIs measured in 2 min time bins and normalized to baseline in response to **C)** quinpirole (<u>WT</u> (black circles), baseline (a): 0.9 ±0.5Hz, 20 min in quinpirole (1): 0.7 ±0.5Hz, N=5, n=15; <u>PNKD</u> (blue squares), baseline (b): 0.9 ±0.3Hz, 20min post-quinpirole (2): 3.4 ±0.4Hz, N=7, n=18), RM 2-way ANOVA, time: F_(9.279)_= 4.08, p<0.0001; genotype: F_(1,31)_= 6.48, p=0.01; interaction: F_(9,279)_= 4.02, p<0.0001; ** at 22–24 min Sidak’s multiple comparisons test; **D)** caffeine (<u>WT</u> (black circles), baseline (a): 1.04 ±0.07Hz, 18min post-caffeine (1): 0.25 ±0.07Hz, N=4, n=11; <u>PNKD</u> (blue squares), baseline (b): 1.12 ±0.05Hz, 18min post-caffeine (2): 0.18 ±0.05Hz, N=6, n=18), RM 2-way ANOVA, time: F_(9,243)_= 92.15, p<0.0001; genotype: F_(1,27)_= 0.94, p=0.34; interaction: F_(9,243)_ =1.044, p=0.4; ns from 4 to 20 min Sidak’s multiple comparisons test and **E)** of quinpirole in the presence of caffeine (<u>WT</u> (black circles), baseline in caffeine (a): 0.52 ±0.2Hz, 12min in quinpirole (1): 1.48 ±0.2Hz, N=4, n=9; <u>PNKD</u> (blue squares), baseline in caffeine (b): 0.34 ±0.2Hz, 12min in quinpirole (2): 0.11 ±0.2Hz, N=3,n=7), RM 2-way ANOVA, time: F_(11,154)_= 2.302, p=0.01; genotype: F_(1,14)_= 6.531, p=0.02; interaction: F_(11,154)_= 3.18, p=0.0006; ** at 22-24min Sidak’s multiple comparisons test. Points are mean + SEM calculated from nested data.

In contrast, in both WT and PNKD mice, caffeine (1µM) induced a pronounced depression of ChI firing rates (**Figure 2D**) with no differences between genotypes.

It is well established that caffeine triggers DA release in the striatum [31]. In the PNKD transgenic mouse, this stimulating effect on DA release has been shown to be proportionally enhanced compared to controls [16]. To mimic the combined effect of caffeine and dopamine D2R activation on ChIs in *ex vivo* electrophysiology experiments, we exposed striatal slices to caffeine and quinpirole sequentially. This combination revealed an unexpected interaction: in the presence of caffeine (1µM), the effects of quinpirole (10µM) in both genotypes were reversed. Indeed, in PNKD mice quinpirole exerted the canonical inhibitory effect on ChI firing rate (**Figure 2E**) but in ChIs from WT mice it induced a paradoxically excitatory effect (**Figure 2E**). Direct comparison between genotypes confirmed that WT mice had significantly higher ChI firing after quinpirole application as compared to PNKD mice, indicating that caffeine can bidirectionally switch the effects of D2R signaling in ChIs, from inhibitory to excitatory in WT mice, and from excitatory to inhibitory in PNKD mice. The same switch from excitatory to inhibitory effects of D2Rs activation was observed by incubating PNKD slices in a selective A2ARs antagonist, ZM241385 (500nM, not shown), suggesting that caffeine’s effects may be mediated by antagonism of A2ARs.

### Manifestation of Dystonia in the PNKD Mouse is Associated with Reduced Striatal Intrinsic ACh Rhythms

To investigate ACh transmission *in vivo*, we monitored real time dynamics of ACh release in striatum using GRAB-ACh-based *in vivo* fiber photometry in awake, behaving PNKD and WT mice (**Figure 3A**). The signal of striatal ACh activity from both genotypes was mainly distributed in the low frequency band (delta frequency: 1.2–2.2 Hz; **Figure 3B-C** [1] and **D**), as previously described [21]. Quantitative analysis showed that the average power of the delta frequency band (% of total sum) was significantly higher in PNKD mice compared to the WT controls at baseline (**Figure 3E**) and was associated with increased baseline locomotion speed (**Figure 3F**), suggesting that higher coherence of ChI population activity in the delta frequencies is associated with hyperlocomotion in PNKD mice at baseline.

**Figure 3.**
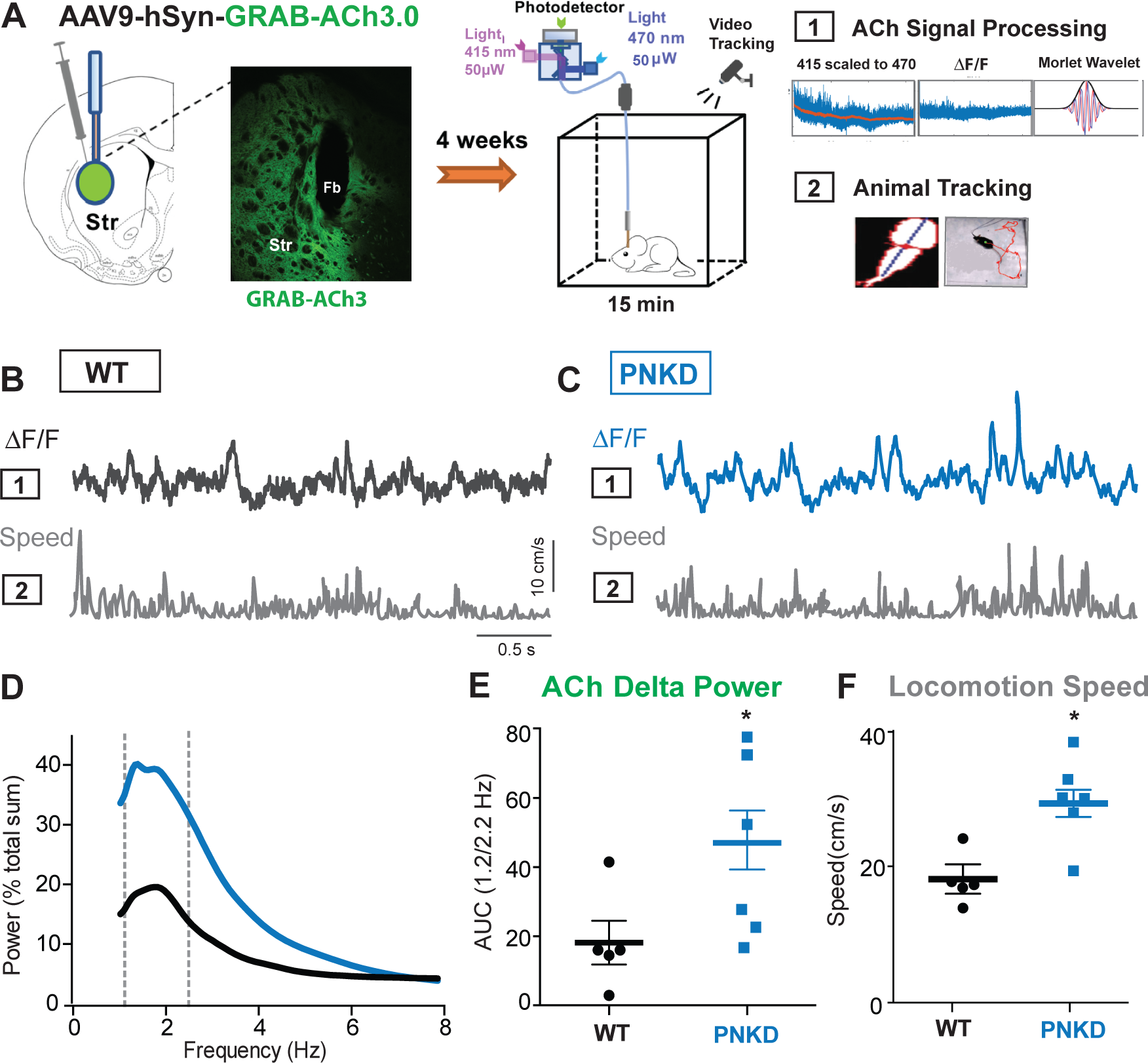
Spontaneous ACh Delta Rhythm and Locomotion Speed are Enhanced in the Pre-Symptomatic PNKD Mouse. **A)** ACh sensor expression in dorsolateral striatum, *in vivo* fiber photometry setup and analysis pipelines. **B)** Example GRAB-ACh fluorescence trace ([1] ΔF/F) in a WT and **C)** in a PNKD mouse during spontaneous movement ([2] instantaneous locomotion speed from animal tracking). **D)** Mean normalized power spectra of the ACh signal at baseline in WT (black, N=5) and PNKD mice (blue, N=6). Vertical dotted lines indicate 1.2-2.2 Hz. **E)** Scatter plots showing area under the curve of the normalized delta band at baseline in WT compared to PNKD mice (Mann-Whitney test, *p=0.026) and **F)** the speed of locomotion at baseline in WT vs PNKD mice (Mann-Whitney test, *p=0.019)

To study the relationship between striatal spontaneous fluctuations of ACh and manifestation of the behavioral phenotype, we treated WT and PNKD mice with caffeine, quinpirole, or saline vehicle (**Figure 4A**). In accordance with the behavioral analysis shown before, we found that treatment with caffeine induced a prominent dystonic phenotype in the majority of PNKD mice (4 out of 6) (**Figure 4D** [1]), whereas only hyperactivity, measured as increased locomotion speed, was observed in WT (**Figure 4C** [3]). Enhanced locomotion speed in WT mice was associated with a sustained increase of the ACh delta band power (**Figure 4B-C**).

**Figure 4.**
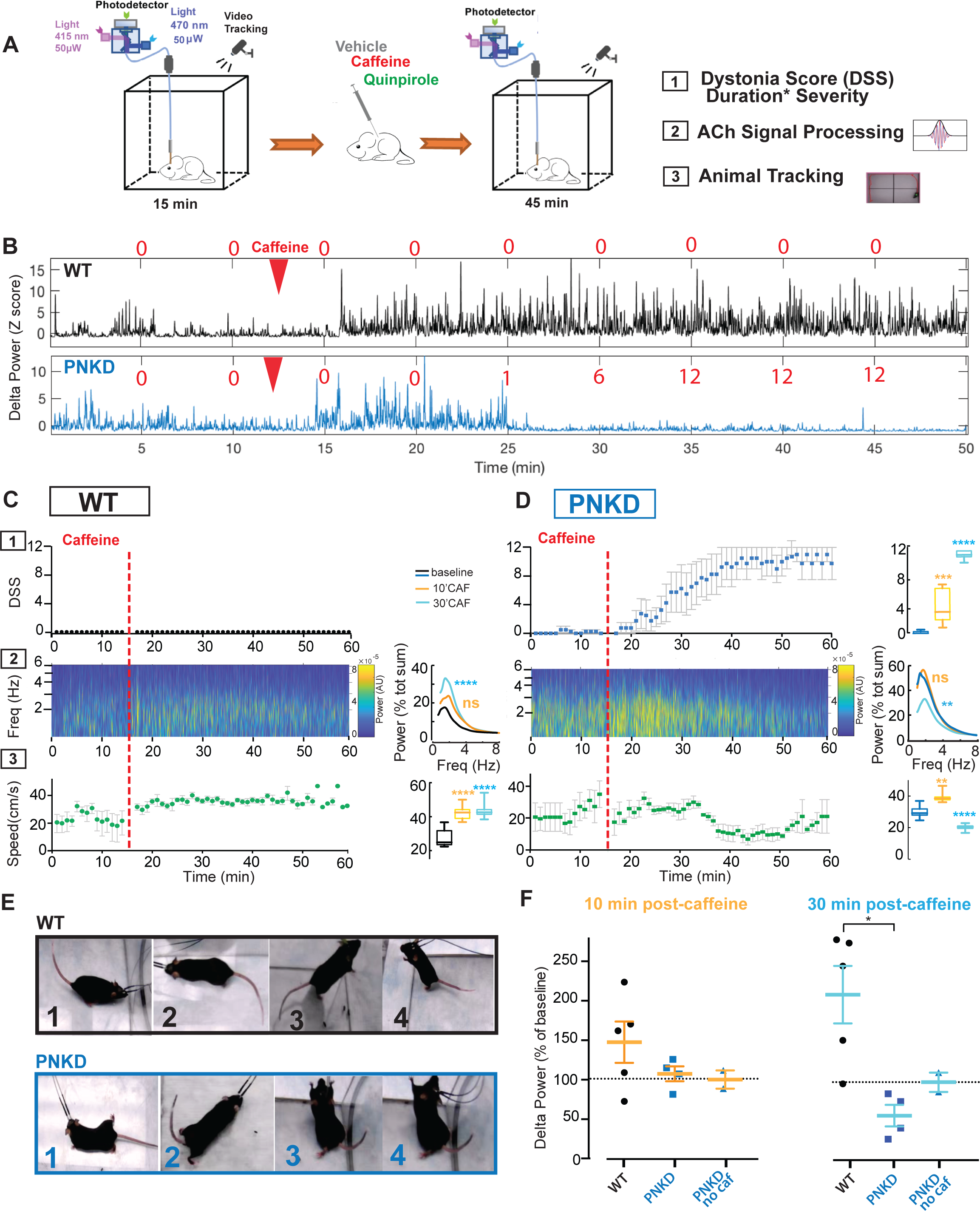
Caffeine-Induced Movement Alterations Correspond to Changes in Spontaneous ACh Delta Rhythm. **A)** *In vivo* fiber photometry experimental design and analysis pipelines. **B)** Representative traces of the ACh signal z-scored to pre-injection baseline in 1WT and 1PNKD mouse before and after caffeine injection (red arrowhead). Red numbers indicate DSS. **C)** WT (N=5) vs **D)** PNKD (N=4) mice time courses and over time summary box plots of: (Top, [1]) average dystonia score obtained with a modified time-based Dystonia Severity Score (DSS). Red dotted line shows caffeine injection at 15 min from the start of recording (PNKD score, RM 1-way ANOVA, treatment: F_(1.084,10.84)_= 211.9, p<0.0001; ***baseline vs 10min; **** baseline vs 30min Dunnett’s multiple comparisons test.) Points are mean + SEM. (Middle, [2]) Colormaps of the average frequency spectra obtained by Morlet Wavelet convolution of the ΔF/F photometry signals. Average over time of the power spectrum shows changes of the delta power at different time windows after caffeine injection (<u>WT</u>: 1-way ANOVA, treatment: F_(2,192)_= 9.2, p=0.0002; ns baseline vs 10min; **** baseline vs 30min, Dunnett’s multiple comparisons test. <u>PNKD</u>: 1-way ANOVA, treatment: F_(2,192)_= 6.5, p=0.0019; ns baseline vs 10min; ** baseline vs 30min, Dunnett’s multiple comparisons test). (Bottom, [3]) Time courses of the locomotion speed and overtime summary box plots (<u>WT</u>: RM 1-way ANOVA, treatment: F_(1.836,18.36)_= 46.9, p<0.0001; **** baseline vs 10min; **** baseline vs 30min Dunnett’s multiple comparisons test. <u>PNKD</u>: RM 1-way ANOVA, treatment: F_(1.907,19.07)_= 107.3, p<0.0001; ** baseline vs 10min; **** baseline vs 30min, Dunnett’s multiple comparisons test). **E)** Photos illustrating the movement phenotype displayed by the mice in B. **F)** Summary scatter plots of caffeine’s effects on ACh delta power in WT vs PNKD mice at two different time points. PNKD treated with vehicle saline (N=2) shown in Supplemental Figure 1 are also included for comparison (1-way ANOVA, genotype/treatment: F_(7,18)_= 3.64, p<0.01, * Tukey’s multiple comparisons test).

Vehicle injections in the PNKD mice only slightly changed the motor behavior since habituated PNKD mice were used in these experiments (**Suppl. Figure 1B** [1] and [3]). Similarly, the ACh delta power did not change over time after vehicle injections **(Figure 4F** and **Suppl. Figure 1B** [2]). In PNKD mice, caffeine increased the speed of locomotion in the first 10 min post-injection prior to the onset of a measurable dystonic phenotype (**Figure 4D** [3]), which was again associated with enhanced ACh delta power (**Figure 4B-D** [2]). As dystonic movements developed and became more severe and long-lasting, i.e. higher dystonia score and lower speed of movement (**Figure 4D** [1]**-**[3]), the ACh delta power gradually decreased to levels below baseline (**Figure 4D** [2]). The percentage change from baseline of the delta power was significantly depressed in the dystonic phase of the PNKD mice (30 min post-injection) compared to WT mice (**Figure 4F**). These findings support a model in which coordinated activation of the ChI population in the delta band promotes movement, while a reduction of the ChI delta rhythm is associated with abnormalities in movement patterns including dystonia.

To test this model we analyzed separately the two PNKD mice that did not develop a dystonic/dyskinetic phenotype in response to caffeine. Similarly to WT mice, these PNKD mice exhibited hyperlocomotion in response to caffeine but not the dystonic/dyskinetic phenotype (**Suppl. Figure 1A** [1] and [3]). In accordance with our hypothetical model, the ACh delta power increased over time as observed in WT mice (**Suppl. Figure 1A** [2]).

To further verify this model, we repeated the photometry experiment using the D2R agonist, quinpirole, which induced locomotor inactivity in both WT and PNKD mice that in some cases lasted the entire testing session. This prolonged inactivity rendered the rating of dystonia unsuitable (**Suppl. Figure 2A-B** [1]). We found that quinpirole-induced immobility was associated with a significant depression of ACh signal power in the delta band in both PNKD and WT (**Suppl. Figure 2A-B** [2]). Overall, these experiments confirmed that the intrinsic ACh delta rhythm changed in accordance with motor activity.

### Changes in Spontaneous ACh Rhythm Correspond to Changes in Striatal ACh Concentration

Fiber photometry records arbitrary fluorescence units without direct indications of objective concentrations of neurotransmitter levels [32] **Figure 5A**). To investigate how spontaneous and drug-induced fluctuations of ACh rhythms recorded by fiber photometry relate to striatal extracellular concentrations of ACh, we performed *in vivo* microdialysis recordings in separate groups of WT and PNKD mice treated with caffeine or quinpirole. As previously observed in the context of dopamine efflux [16], we found that basal extracellular ACh was significantly reduced in PNKD compared to WT littermates (**Figure 5B**).

**Figure 5.**
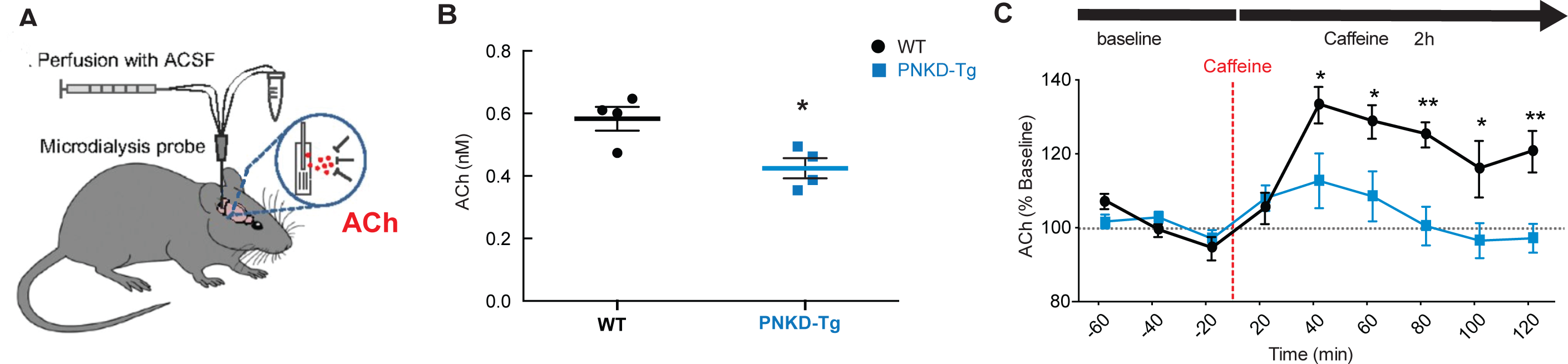
Caffeine Reduced Striatal Extracellular ACh Levels in PNKD Mice. **A)** *In vivo* microdialysis setup. **B)** Extracellular ACh concentration is reduced in PNKD vs WT mice at baseline (Unpaired t-test, t(6)=3.183, p=0.0190). **C)** Time course of the % change of ACh concentration in WT vs PNKD mice before and after caffeine injections (RM 2-way ANOVA, time: F_(8,112)=_9.555, p<0.0001; genotype:F_(1,14)=_6.860, p=0.0202; time x genotype:F_(8,112)=_4.860, p<0.0001. Bonferroni post-hoc comparisons from 40 min to 120 min, all p<0.05). Points are mean±SEM.

In order to determine whether the disruption of the ACh rhythm associated with the mixed dystonia/dyskinesia attacks correspond to reduced striatal ACh efflux, we assessed ACh levels upon caffeine administration during *in vivo* microdialysis. We observed an increase in ACh in both WT and PNKD mice beginning 40 min after caffeine injection (**Figure 5C**, approximately 20%), closely associated with the onset of abnormal movements in PNKD and hyperactivity in WT mice. However, caffeine-induced ACh efflux was attenuated compared to WT during this same time period. At time points between 80 and 120 min, ACh levels actually dipped below those observed at baseline. These time points were associated with a prominent and stable motor phenotype. Similarly, quinpirole (1mg/kg) induced depression of the ACh levels associated with immobility at both time points (**Suppl. Figure 2C**).

These results show that, with the expected differences in time resolution, the overall time course of caffeine-induced ACh release correlates well between microdialysis and fiber photometry, with the peak of ACh release within the first 20 min from injection in both genotypes and a faster decay in PNKD compared to WT mice.

## Discussion

In this study we have investigated the role of striatal ChIs in the development of hyperkinetic movements in an inducible phenotypic transgenic mouse model of paroxysmal non-kinesigenic dyskinesia/dystonia (PNKD). In the course of this work, we have extended the characterization of the PNKD mouse model, demonstrating the presence of prominent inducible dystonic features. We have found that there is a close relationship between the appearance of dystonic movements and alterations in cholinergic transmission. These observations shed new light on the common finding of D2R-induced ChI “paradoxical excitation” in models of dystonia, suggesting this is a consequence of dynamic regulation of ChI function.

### Behavioral Characterization of the Inducible Motor Phenotype in the PNKD Mice

Humans carrying the PNKD gene mutation develop attacks of dystonic posturing with some choreic and ballistic movements, often precipitated by caffeine. Similarly, PNKD transgenic mice develop a hyperkinetic motor disorder upon exposure to caffeine. The inducible phenotype observed in the PNKD mice has been mainly described as dyskinesia in prior literature [16, 18]. Indeed, we did observe dyskinesia, which was predominantly orofacial, but we also observed prominent and sustained dystonic features in the mouse model. The postural abnormalities observed in the PNKD animals treated with caffeine are very similar to previous descriptions of rodent dystonia induced by DA antagonists or cerebellar dysfunction, i.e. twisting of the limbs, hyperextension of tail and digits, and hyperflexion of the neck and back muscles [29, 33]. The manifestation of motor symptoms akin to human dystonia in the PNKD mice in response to external stimuli provides a robust platform for studying the mechanistic basis of dystonic movements. Furthermore, the consistent and predictable observation of dystonia-like symptoms in this model bridges the gap left by previous genetic dystonia models lacking a dystonic motor phenotype, offering a novel avenue for exploring the pathophysiology of hyperkinetic conditions.

### D2R “Paradoxical Excitation” of ChIs

In this study we found that PNKD mice exhibit the so-called “paradoxical excitation” of striatal ChIs firing upon D2R activation with quinpirole in *ex vivo* brain slice recordings, a shared endophenotype observed in other non-phenotypic genetic dystonia models. A key finding here was that quinpirole’s excitatory effect was reversed in the presence of the dystonia-inducing drug caffeine *ex vivo*. It is probable that this reversal was mediated by antagonism of A2ARs, since selectively blocking A2ARs mimicked the effect of caffeine, similarly to our previous findings in the *Gnal+/-* genetic mouse model of dystonia [11]. However, we have previously shown that the excitatory response of ChIs to D2Rs activation is not exclusive to disease models, but can be induced in WT mice upon activation of muscarinic receptors in *ex vivo* slices [10]. Similarly, here we found that in WT mice the quinpirole response of ChIs switched to excitatory in the presence of caffeine, which *in vivo* stimulated movement without inducing movement abnormalities. Therefore caffeine, likely through antagonism of A2ARs, can interfere with the ongoing signaling of D2Rs and bidirectionally reverse its effects on ChI firing rate. These findings challenge the conventional understanding of the D2R-mediated inhibition of ChIs and point towards a more complex and dynamic interplay between D2Rs and other GPCRs in shaping the D2R signal transduction and downstream ChI activity. There is a vast literature describing D2R functional heteromers with a variety of other GPCRs, which increase both the complexity and diversity of receptor-mediated signal transduction. In functional heteromers, the downstream signaling dynamically switches based on the activity level and also the dynamics (i.e. the order in which GPCRs are activated) of participating neurotransmitters, resulting in disparate consequences on cellular physiology and function [34–39]. Together these observations challenge the idea of D2R excitatory signaling as “paradoxical” and point toward a new interpretation in which D2R signaling is dynamically regulated by complex receptor-receptor interactions. These may culminate to drive compensatory and/or protective mechanism(s) in response to certain stimuli (caffeine) or insults (*PNKD* or other gene mutations associated with dystonia), preventing manifestation of movement abnormalities. Ultimately, these findings are important to discern the causative versus compensatory nature of the ChI “paradoxical excitation” previously described in non-phenotypic genetic models.

### ACh Delta Rhythm and Movement

Recent work established a correlation between the magnitude of ACh fluctuations and the degree of spike coherence among ChIs [21]. By recording the discharge of ChIs while simultaneously imaging ACh in overlapping regions of dorsal striatum, the study demonstrated that coherent ChI firing drives periodic ACh fluctuations mainly distributed in the delta frequency, the amplitude of which reflects the degree of local spike coherence (synchronously active neurons) among neighboring ChIs. Furthermore, an independent study reported that delta-frequency patterned dynamics of individual ChIs are coupled to animals’ stepping cycles, suggesting that this delta rhythmicity plays an important role in patterning of movement [22].

In photometry recordings of ACh dynamics in awake behaving animals, we found that in WT mice caffeine’s stimulatory effect on locomotion was associated with enhanced power of striatal ACh delta rhythm. On the contrary, caffeine-induced movement abnormalities resembling dystonic symptoms in the PNKD mice, as well as quinpirole-induced depression of voluntary movement in both genotypes, coincided with sustained depression of the striatal ACh delta rhythm. Disruptions of this rhythm were specifically associated with manifestation of movement abnormalities since the delta power was not reduced upon vehicle injections in PNKD animals or in mice carrying the PNKD mutation that did not develop a motor phenotype in response to caffeine.

Overall, these findings support the notion that ACh oscillations contribute to regulation of motor activity. Enhanced synchronization among ChIs may underlie the heightened ACh delta power detected in pre-symptomatic PNKD and caffeine-treated WT mice, which could play a role in orchestrating the enhanced mobility. On the other hand, loss of coherence in the ChI network could underlie the depression of ACh oscillations leading to disruption of natural movement sequencing typical of dystonic symptoms.

Microdialysis investigations also provided insights into how changes in ACh delta oscillations are reflected in extracellular levels of striatal ACh. We found that caffeine’s effects on ACh concentration mirrored the effects on the power of ACh delta oscillations detected by fiber photometry, with the expected delay from the time of injection due to different temporal resolution between the techniques, suggesting that coordinated activation of ChIs is reflected in enhanced tonic levels of ACh. However, despite high translatability between microdialysis and photometry in response to caffeine challenge, ACh levels at baseline do not seems to reflect the magnitude of ACh delta oscillations. Indeed, PNKD mice showed enhanced ACh delta power prior to exposure to caffeine compared to WT mice, but decreased levels of extracellular ACh at baseline. This discrepancy is possibly due to either higher variance of microdialysis measures across subjects or spatial differences (i.e. diffusion to the probe in microdialysis vs fluorescence generated in microdomains at immediate site of release).

### Do D2Rs Modulate Movement Through Regulation of the ACh Delta Rhythm?

Although other studies have indicated that ChIs contribute to the regulation of motor activity upon D2R stimulation [13, 14, 40], this is the first study associating motor abnormalities to changes in spontaneous ACh delta rhythm and to changes in D2R modulation of ChI activity.

Here, we have found that conditions that resulted in enhanced locomotor activity and increased power of ACh delta oscillations *in vivo* correspond to excitatory D2R effects on ChI activity in *ex vivo.* In addition, caffeine can disrupt spontaneous movement and ACh delta rhythm in PNKD behaving animals while at the cellular level abolishing the D2R excitation of ChIs. These observations lead us speculate a role for excitatory D2R signaling in synchronizing firing across ensembles of ChIs to produce ACh delta oscillations that allow for proper sequencing and patterning of movements (Figure 6). Indeed, blockade of D2Rs with first generation antipsychotics induces debilitating side effects known as extrapyramidal symptoms, which include dystonia, parkinsonism, and tardive dyskinesia [14]. Future studies to directly investigate how D2Rs on ChIs regulate their spontaneous delta rhythm in different activity states will require *in vivo* monitoring of ChI oscillations in mice with cell-type-specific ablation of D2R in ChIs. Moreover, the use of functionally biased D2R ligands [41, 42–44] could be advantageous to segregate the behavioral effects of distinct signaling pathways.

**Figure 6.**
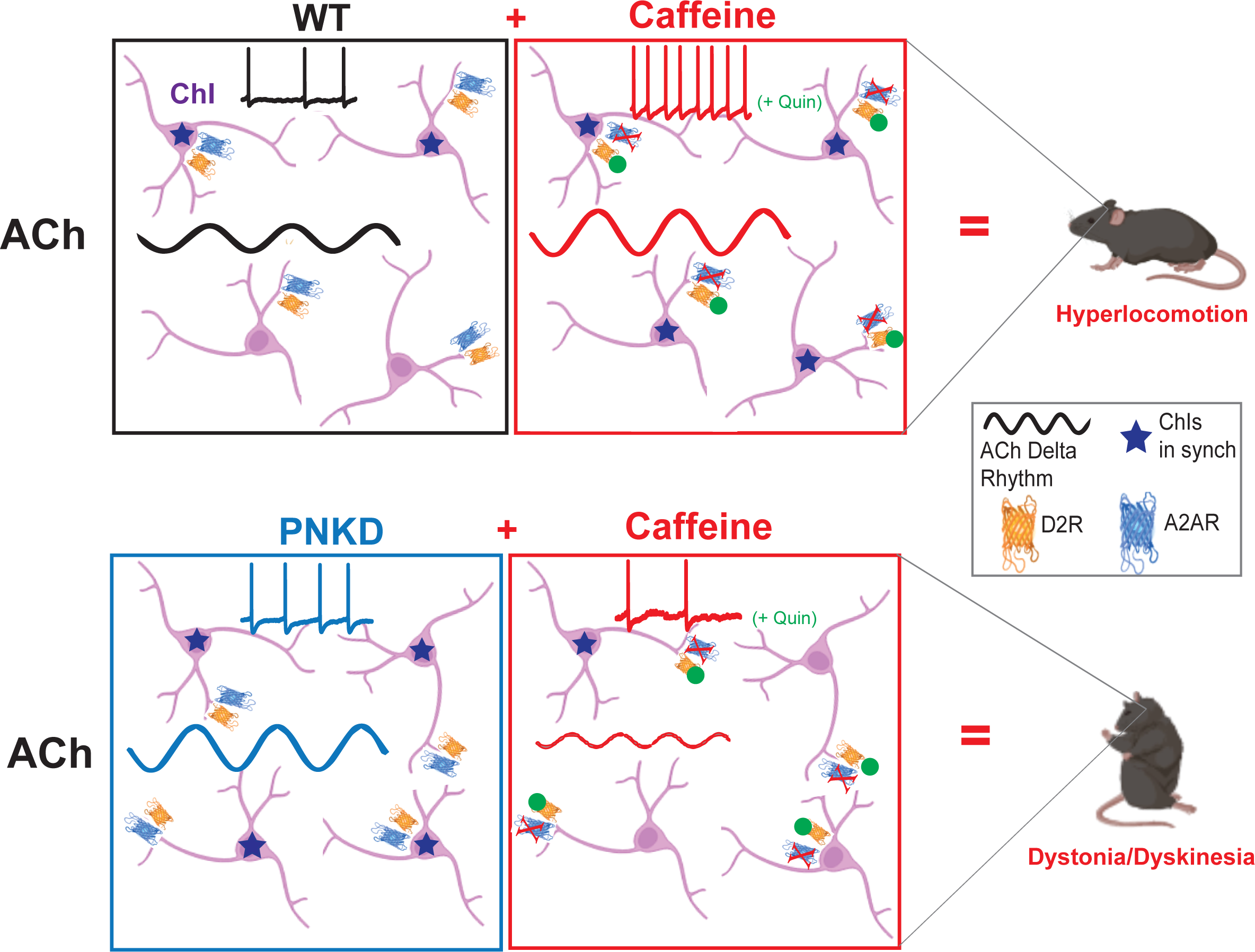
Summary Working Model. Shows a network of striatal ChIs which can synchronize their spontaneous firing activity to produce ACh oscillations in the delta frequency. In WT mice exposed to caffeine, the amplitude of the ACh delta rhythm increases (more synchrony among ChIs) as the animals become hyperactive and the effects of D2Rs activation on ChI discharge become excitatory at the cellular level (Top right). This behavioral, cellular and network profile is mirrored by pre-symptomatic PNKD mice before caffeine treatment (Bottom left). Upon exposure to caffeine, PNKD mice develop a motor phenotype with dyskinesia and dystonia features associated with reduced ACh delta oscillations and loss of D2Rs excitatory effects on ChIs firing (Bottom right).

### Conclusions

In summary, our study demonstrates a direct correlation between striatal ACh delta rhythm and motor regulation. Caffeine’s stimulatory effect on locomotion in WT mice is linked to increased ACh delta power, while caffeine induces dystonic-like symptoms in PNKD mice, accompanied by sustained depression of ACh delta rhythm. The disruption of ACh oscillations specifically aligns with movement abnormalities, emphasizing the pivotal role of ACh dynamics in motor control.

Our findings also highlight the connection between excitatory dopamine D2R effects on ChI activity and enhanced ACh delta oscillations, suggesting an essential role for excitatory D2R signaling in coordinating ChI firing for the generation of ACh delta oscillations crucial for proper movement sequencing. In this view, the phenomenon of “paradoxical excitation” of ChIs described *ex vivo* in several non-phenotipic genetic mouse models of dystonia can be interpreted as a compensatory or protective mechanism that prevents manifestation of movement abnormalities.

Overall, our study unveils the intricate interplay between ACh dynamics, ChI activity, and motor regulation, providing insights for future research and potential therapeutic interventions in movement disorders.

---

**Suppl. Figure 1.**
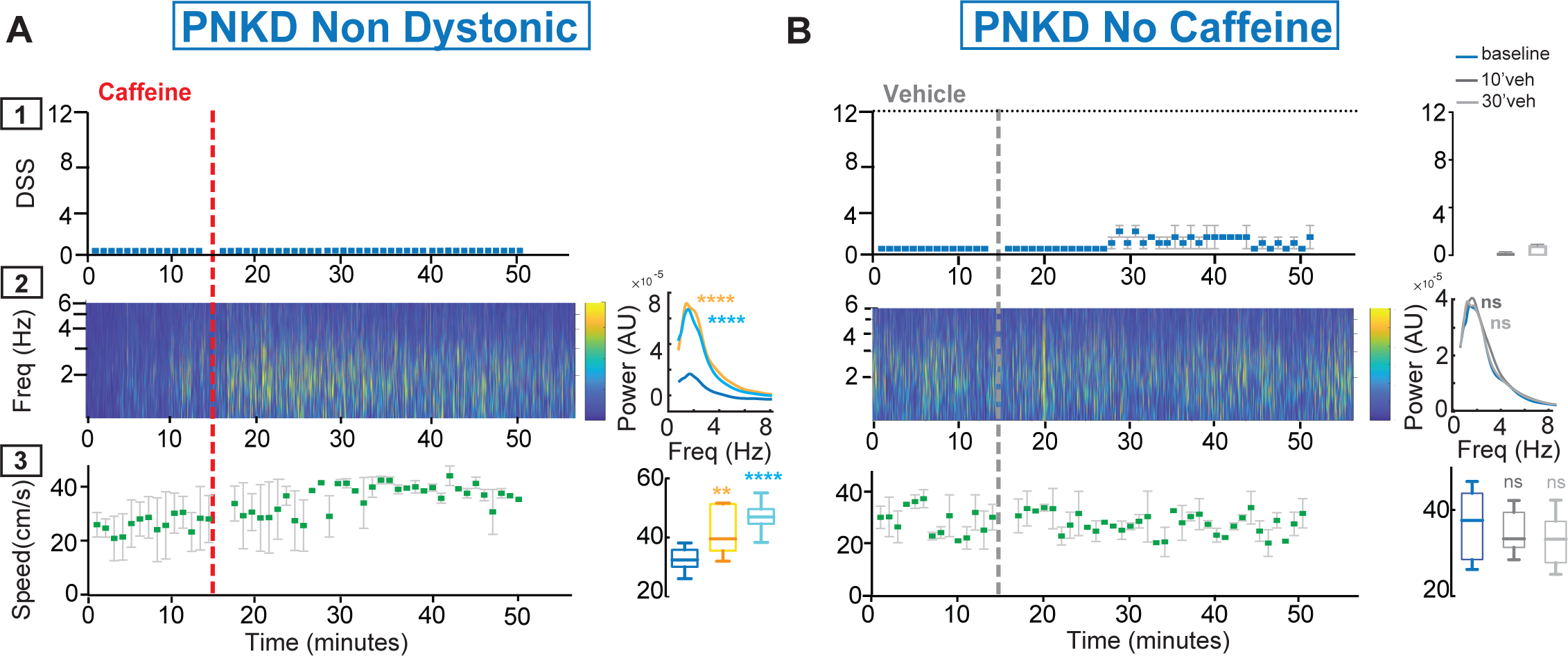
The ACh Delta Rhythm Is Not Reduced in the Absence of Motor Phenotype in PNKD Mice. **A)** (Top) Time courses of the average Dystonia Severity Score (DSS) induced by injection of caffeine (red dotted line) in non-phenotypic PNKD mice (N=2). (Middle) Color plots of the average frequency spectrum obtained by Morlet Wavelet convolution of the ΔF/F photometry signals. Average over time of the power spectrum shows changes of the delta power at different time windows after caffeine (1-way ANOVA, treatment: F_(2,192)_= 26.5, p<0.0001, **** baseline vs 10 min and **** baseline vs 30 min, Dunnett’s multiple comparisons test). (Bottom) Time courses of the movement speed and overtime summary bar charts (RM 1-way ANOVA, treatment: F_(1.521,15.21)_= 16.4, p=0.0003, ** baseline vs 10 min and **** baseline vs 30 min, Dunnett’s multiple comparisons test). **B)** (Top) Time courses of the average Dystonia Severity Score (DSS) induced by injection of saline vehicle (gray dotted line) in the same PNKD mice shown in Fig. 3 and 4 (RM 1-way ANOVA, treatment: F_(1.078,10.78)_= 24.1, p=0.0004, ns baseline vs 10 min and 30 min, Dunnett’s multiple comparisons test). (Middle) Color plots of the average frequency spectrum and average over time bar charts show no changes in delta power after saline injections (1-way ANOVA, treatment: F_(2,192)_= 0.27, p=0.8, ns baseline vs 10 min and ns baseline vs 30 min, Dunnett’s multiple comparisons test). (Bottom) Time courses of the movement speed and overtime summary bar charts (RM 1-way ANOVA, treatment: F_(1.818,18.18)_= 0.86, p=0.43, ns baseline vs 10 min and ns baseline vs 30 min, Dunnett’s multiple comparisons test).

**Suppl. Figure 2.**
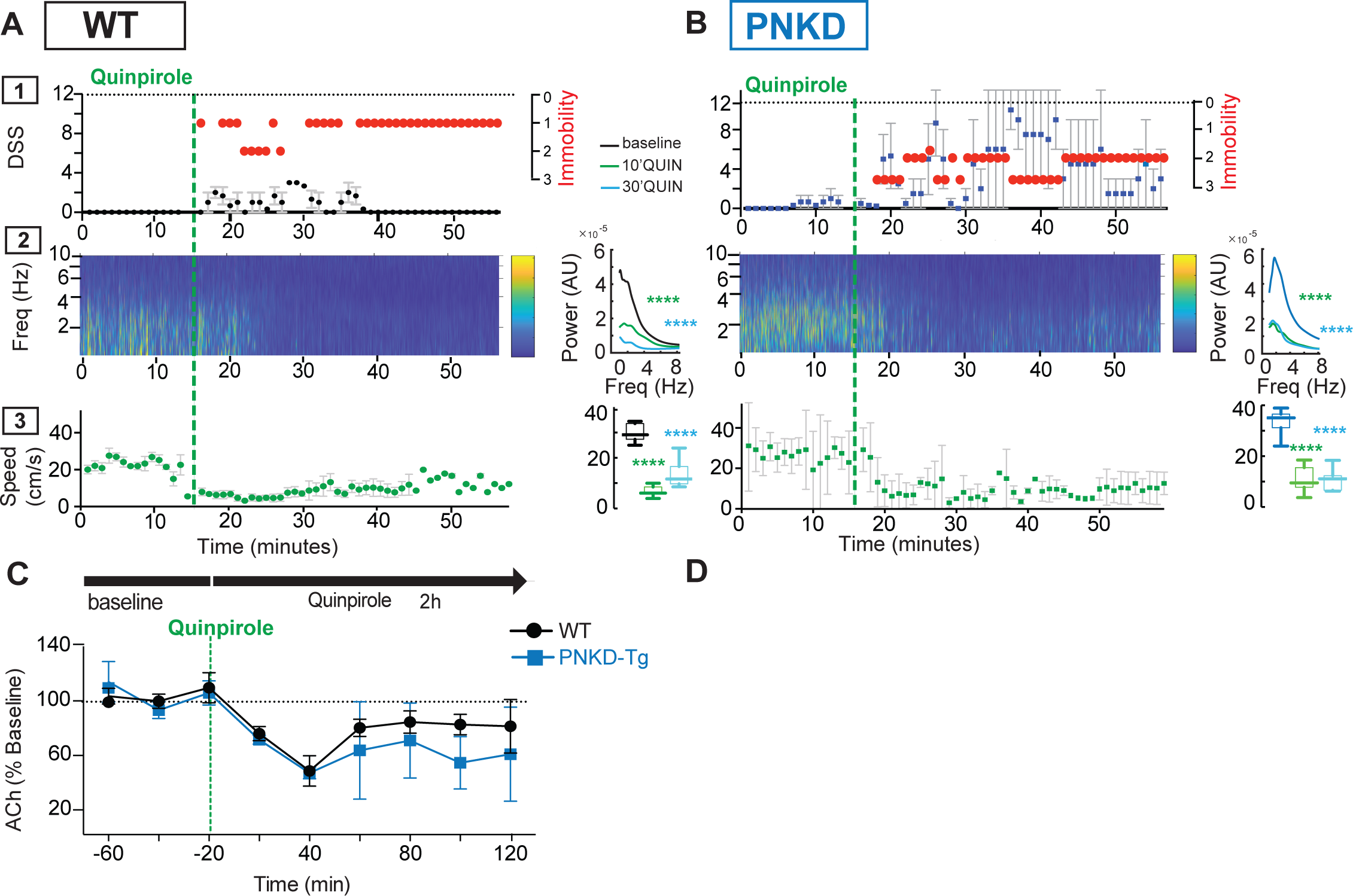
Quinpirole-Induced Immobility is Associated with Disrupted ACh Delta Rhythms. **A)** WT (N=4) and **B)** PNKD (N=4) mice (Top) Time courses of the average Dystonia Severity Score (DSS) + (red dots) immobility score induced by injection of quinpirole (green dotted line). (Middle) Color plots of the average frequency spectra of the ACh photometry signals. Average over time of the power spectrum shows depression of the delta power at all time windows after quinpirole injection (<u>WT</u>: 1-way ANOVA, treatment: F_(2,190)_= 55.2, p<0.0001, ****baseline vs 10 min and **** baseline vs 30 min, Dunnett’s multiple comparisons test; <u>PNKD</u>: 1-way ANOVA, treatment: F_(2,192)_= 66.8, p<0.0001, **** baseline vs 10 min and **** baseline vs 30 min, Dunnett’s multiple comparisons test) with no differences between genotypes (2-way ANOVA, time: F_(2,256)_= 253.9, p=0.0001; genotype: F_(1,128)_= 5.82, p=0.01; interaction: F_(2,256)_ =9.337, p=0.0001, *at baseline, ns at 10min, *at 10 min, Sidak’s multiple comparisons test). (Bottom) Time courses of the movement speed and overtime summary bar charts (<u>WT:</u> RM 1-way ANOVA, treatment: F_(1.413,14.13)_= 154.2, p<0.0001, **** baseline vs 10 min and **** baseline vs 30 min, Dunnett’s multiple comparisons test; <u>PNKD:</u> RM 1-way ANOVA, treatment: F_(1.943,19.43)_= 122.3, p<0.0001, **** baseline vs 10 min and **** baseline vs 30 min, Dunnett’s multiple comparisons test). **C)** Time course of the % change of ACh concentration in WT vs PNKD mice before and after quinpirole injections (RM 2-way ANOVA, time: F_(8,16)=_2.1, p=0.09; genotype: F_(1,2)=_0.13, p=0.76; time x genotype: F_(8,16)=_0.59, p=0.77). Points are mean±SEM.

**Suppl. Video 1. Dystonic and Dyskinetic Features of the PNKD Motor Phenotype.** The video shows a WT mouse rendered hyperactive by treatment with 20 mg/Kg of caffeine compared to the motor response of a PNKD mouse, which displays movement abnormalities with dystonic and dyskinetic features.

## References

1. Medina, A., C. Nilles, D. Martino, C. Pelletier, and T. Pringsheim, The Prevalence of Idiopathic or Inherited Isolated Dystonia: A Systematic Review and Meta-Analysis. Mov Disord Clin Pract, 2022. 9(7): p. 860–868.

2. Di Fonzo, A., H.A. Jinnah, and M. Zech, Dystonia genes and their biological pathways. Int Rev Neurobiol, 2023. 169: p. 61–103.

3. Vidailhet, M., A. Méneret, and E. Roze, Dystonia: genetics, phenomenology, and pathophysiology. Lancet Neurol, 2020. 19(11): p. 881–882.

4. Neychev, V.K., R.E. Gross, S. Lehéricy, E.J. Hess, and H.A. Jinnah, The functional neuroanatomy of dystonia. Neurobiol Dis, 2011. 42(2): p. 185–201.

5. Standaert, D.G., Update on the pathology of dystonia. Neurobiol Dis, 2011. 42(2): p. 148–51.

6. Goodchild, R.E., K. Grundmann, and A. Pisani, New genetic insights highlight ‘old’ ideas on motor dysfunction in dystonia. Trends Neurosci, 2013. 36(12): p. 717–25.

7. Howe, M., I. Ridouh, A.L. Allegra Mascaro, A. Larios, M. Azcorra, and D.A. Dombeck, Coordination of rapid cholinergic and dopaminergic signaling in striatum during spontaneous movement. Elife, 2019. 8.

8. Eskow Jaunarajs, K.L., P. Bonsi, M.F. Chesselet, D.G. Standaert, and A. Pisani, Striatal cholinergic dysfunction as a unifying theme in the pathophysiology of dystonia. Prog Neurobiol, 2015. 127-128: p. 91-107.

9. Stahl, S.M., K.L. Davis, and P.A. Berger, The neuropharmacology of tardive dyskinesia, spontaneous dyskinesia, and other dystonias. J Clin Psychopharmacol, 1982. 2(5): p. 321–8.

10. Scarduzio, M., C.N. Zimmerman, K.L. Jaunarajs, Q. Wang, D.G. Standaert, and L.L. McMahon, Strength of cholinergic tone dictates the polarity of dopamine D2 receptor modulation of striatal cholinergic interneuron excitability in DYT1 dystonia. Exp Neurol, 2017. 295: p. 162–175.

11. Eskow Jaunarajs, K.L., M. Scarduzio, M.E. Ehrlich, L.L. McMahon, and D.G. Standaert, Diverse Mechanisms Lead to Common Dysfunction of Striatal Cholinergic Interneurons in Distinct Genetic Mouse Models of Dystonia. J Neurosci, 2019. 39(36): p. 7195–7205.

12. Pisani, A., G. Martella, A. Tscherter, P. Bonsi, N. Sharma, G. Bernardi, and D.G. Standaert, Altered responses to dopaminergic D2 receptor activation and N-type calcium currents in striatal cholinergic interneurons in a mouse model of DYT1 dystonia. Neurobiol Dis, 2006. 24(2): p. 318–25.

13. Lewis, R.G., M. Serra, D. Radl, M. Gori, C. Tran, S.E. Michalak, C.D. Vanderwal, and E. Borrelli, Dopaminergic Control of Striatal Cholinergic Interneurons Underlies Cocaine-Induced Psychostimulation. Cell Rep, 2020. 31(3): p. 107527.

14. Kharkwal, G., K. Brami-Cherrier, J.E. Lizardi-Ortiz, A.B. Nelson, M. Ramos, D. Del Barrio, D. Sulzer, A.C. Kreitzer, and E. Borrelli, Parkinsonism Driven by Antipsychotics Originates from Dopaminergic Control of Striatal Cholinergic Interneurons. Neuron, 2016. 91(1): p. 67–78.

15. Augustin, S.M., J.H. Chancey, and D.M. Lovinger, Dual Dopaminergic Regulation of Corticostriatal Plasticity by Cholinergic Interneurons and Indirect Pathway Medium Spiny Neurons. Cell Rep, 2018. 24(11): p. 2883–2893.

16. Lee, H.Y., J. Nakayama, Y. Xu, X. Fan, M. Karouani, Y. Shen, E.N. Pothos, E.J. Hess, Y.H. Fu, R.H. Edwards, and L.J. Ptácek, Dopamine dysregulation in a mouse model of paroxysmal nonkinesigenic dyskinesia. J Clin Invest, 2012. 122(2): p. 507–18.

17. Scarduzio, M. and D.G. Standaert, Piecing together a complex puzzle: 5 key challenges in basic dystonia research. Dystonia, 2023. 2.

18. Nelson, A.B., A.E. Girasole, H.Y. Lee, L.J. Ptáček, and A.C. Kreitzer, Striatal Indirect Pathway Dysfunction Underlies Motor Deficits in a Mouse Model of Paroxysmal Dyskinesia. J Neurosci, 2022. 42(13): p. 2835–2848.

19. Shen, Y., W.P. Ge, Y. Li, A. Hirano, H.Y. Lee, A. Rohlmann, M. Missler, R.W. Tsien, L.Y. Jan, Y.H. Fu, and L.J. Ptáček, Protein mutated in paroxysmal dyskinesia interacts with the active zone protein RIM and suppresses synaptic vesicle exocytosis. Proc Natl Acad Sci U S A, 2015. 112(10): p. 2935–41.

20. Erro, R., Familial Paroxysmal Nonkinesigenic Dyskinesia, in GeneReviews(®), M.P. Adam, et al., Editors. 1993, © 1993-2024, University of Washington, Seattle. GeneReviews is a registered trademark of the University of Washington, Seattle: Seattle WA.

21. Krok, A.C., M. Maltese, P. Mistry, X. Miao, Y. Li, and N.X. Tritsch, Intrinsic dopamine and acetylcholine dynamics in the striatum of mice. Nature, 2023. 621(7979): p. 543-549.

22. Shroff, S.N., E. Lowet, S. Sridhar, H.J. Gritton, M. Abumuaileq, H.A. Tseng, C. Cheung, S.L. Zhou, K. Kondabolu, and X. Han, Striatal cholinergic interneuron membrane voltage tracks locomotor rhythms in mice. Nat Commun, 2023. 14(1): p. 3802.

23. Lundblad, M., A. Usiello, M. Carta, K. Håkansson, G. Fisone, and M.A. Cenci, Pharmacological validation of a mouse model of l-DOPA-induced dyskinesia. Exp Neurol, 2005. 194(1): p. 66–75.

24. Jinnah, H.A., J.P. Sepkuty, T. Ho, S. Yitta, T. Drew, J.D. Rothstein, and E.J. Hess, Calcium channel agonists and dystonia in the mouse. Mov Disord, 2000. 15(3): p. 542–51.

25. Pizoli, C.E., H.A. Jinnah, M.L. Billingsley, and E.J. Hess, Abnormal cerebellar signaling induces dystonia in mice. J Neurosci, 2002. 22(17): p. 7825–33.

26. Raike, R.S., C.E. Pizoli, C. Weisz, A.M. van den Maagdenberg, H.A. Jinnah, and E.J. Hess, Limited regional cerebellar dysfunction induces focal dystonia in mice. Neurobiol Dis, 2013. 49: p. 200–10.

27. Rose, S.J., X.Y. Yu, A.K. Heinzer, P. Harrast, X. Fan, R.S. Raike, V.B. Thompson, J.F. Pare, D. Weinshenker, Y. Smith, H.A. Jinnah, and E.J. Hess, A new knock-in mouse model of l-DOPA-responsive dystonia. Brain, 2015. 138(Pt 10): p. 2987–3002.

28. Jing, M., Y. Li, J. Zeng, P. Huang, M. Skirzewski, O. Kljakic, W. Peng, T. Qian, K. Tan, J. Zou, S. Trinh, R. Wu, S. Zhang, S. Pan, S.A. Hires, M. Xu, H. Li, L.M. Saksida, V.F. Prado, T.J. Bussey, M.A.M. Prado, L. Chen, H. Cheng, and Y. Li, An optimized acetylcholine sensor for monitoring in vivo cholinergic activity. Nat Methods, 2020. 17(11): p. 1139–1146.

29. Andreoli, L., M. Abbaszadeh, X. Cao, and M.A. Cenci, Distinct patterns of dyskinetic and dystonic features following D1 or D2 receptor stimulation in a mouse model of parkinsonism. Neurobiol Dis, 2021. 157: p. 105429.

30. Feldmann, L.K., R. Lofredi, W.J. Neumann, B. Al-Fatly, J. Roediger, B.H. Bahners, P. Nikolov, T. Denison, A. Saryyeva, J.K. Krauss, K. Faust, E. Florin, A. Schnitzler, G.H. Schneider, and A.A. Kühn, Toward therapeutic electrophysiology: beta-band suppression as a biomarker in chronic local field potential recordings. NPJ Parkinsons Dis, 2022. 8(1): p. 44.

31. Negida, A., M. Elfil, Attia, E. Farahat, M. Gabr, A. Essam, D. Attia, and H. Ahmed, Caffeine; the Forgotten Potential for Parkinson’s Disease. CNS Neurol Disord Drug Targets, 2017. 16(6): p. 652–657.

32. Ejdrup, A.L., J. Wellbourne-Wood, J.K. Dreyer, N. Guldhammer, M.D. Lycas, U. Gether, B.J. Hall, and G. Sørensen, Within-Mice Comparison of Microdialysis and Fiber Photometry-Recorded Dopamine Biosensor during Amphetamine Response. ACS Chem Neurosci, 2023. 14(9): p. 1622–1630.

33. Brown, A.M., M.E. van der Heijden, H.A. Jinnah, and R.V. Sillitoe, Cerebellar Dysfunction as a Source of Dystonic Phenotypes in Mice. Cerebellum, 2023. 22(4): p. 719–729.

34. Guidolin, D., C. Tortorella, M. Marcoli, C. Cervetto, R. De Caro, G. Maura, and L.F. Agnati, Modulation of Neuron and Astrocyte Dopamine Receptors via Receptor-Receptor Interactions. Pharmaceuticals (Basel), 2023. 16(10).

35. Fuxe, K., A. Tarakanov, W. Romero Fernandez, L. Ferraro, S. Tanganelli, M. Filip, L.F. Agnati, P. Garriga, Z. Diaz-Cabiale, and D.O. Borroto-Escuela, Diversity and Bias through Receptor-Receptor Interactions in GPCR Heteroreceptor Complexes. Focus on Examples from Dopamine D2 Receptor Heteromerization. Front Endocrinol (Lausanne), 2014. 5: p. 71.

36. Borroto-Escuela, D.O., A.O. Tarakanov, D. Guidolin, F. Ciruela, L.F. Agnati, and K. Fuxe, Moonlighting characteristics of G protein-coupled receptors: focus on receptor heteromers and relevance for neurodegeneration. IUBMB Life, 2011. 63(7): p. 463–72.

37. Pollack, A., Coactivation of D1 and D2 dopamine receptors: in marriage, a case of his, hers, and theirs. Sci STKE, 2004. 2004(255): p. pe50.

38. Rendón-Ochoa, E.A., M. Padilla-Orozco, V.M. Calderon, V.H. Avilés-Rosas, O. Hernández-González, T. Hernández-Flores, M.B. Perez-Ramirez, M. Palomero-Rivero, E. Galarraga, and J. Bargas, Dopamine D(2) and Adenosine A(2A) Receptors Interaction on Ca(2+) Current Modulation in a Rodent Model of Parkinsonism. ASN Neuro, 2022. 14: p. 17590914221102075.

39. Nagaoka, K., N. Asaoka, K. Nagayasu, H. Shirakawa, and S. Kaneko, Enhancement of adenosine A(2A) signaling improves dopamine D(2) receptor antagonist-induced dyskinesia via β-arrestin signaling. Front Neurosci, 2022. 16: p. 1082375.

40. Chancey, J.H., C. Kellendonk, J.A. Javitch, and D.M. Lovinger, Dopaminergic D2 receptor modulation of striatal cholinergic interneurons contributes to sequence learning. bioRxiv, 2023.

41. Sahlholm, K., M. Gómez-Soler, M. Valle-León, M. López-Cano, J.J. Taura, F. Ciruela, and V. Fernández-Dueñas, Antipsychotic-Like Efficacy of Dopamine D(2) Receptor-Biased Ligands is Dependent on Adenosine A(2A) Receptor Expression. Mol Neurobiol, 2018. 55(6): p. 4952–4958.

42. Bonifazi, A., H. Yano, A.M. Guerrero, V. Kumar, A.F. Hoffman, C.R. Lupica, L. Shi, and A.H. Newman, Novel and Potent Dopamine D(2) Receptor Go-Protein Biased Agonists. ACS Pharmacol Transl Sci, 2019. 2(1): p. 52–65.

43. Allen, J.A., J.M. Yost, V. Setola, X. Chen, M.F. Sassano, M. Chen, S. Peterson, P.N. Yadav, X.P. Huang, B. Feng, N.H. Jensen, X. Che, X. Bai, S.V. Frye, W.C. Wetsel, M.G. Caron, J.A. Javitch, B.L. Roth, and J. Jin, Discovery of β-arrestin-biased dopamine D2 ligands for probing signal transduction pathways essential for antipsychotic efficacy. Proc Natl Acad Sci U S A, 2011. 108(45): p. 18488–93.

44. Männel, B., M. Jaiteh, A. Zeifman, A. Randakova, D. Möller, H. Hübner, P. Gmeiner, and J. Carlsson, Structure-Guided Screening for Functionally Selective D(2) Dopamine Receptor Ligands from a Virtual Chemical Library. ACS Chem Biol, 2017. 12(10): p. 2652–2661.

